# Experience-related remapping of temporal encoding by striatal ensembles

**DOI:** 10.1101/2021.03.12.435177

**Authors:** R. Austin. Bruce, Matthew A. Weber, Rachael A. Volkman, Mayu Oya, Eric B. Emmons, Youngcho Kim, Nandakumar S. Narayanan

## Abstract

Temporal control of action is key for a broad range of behaviors and is disrupted in human diseases such as Parkinson’s disease and schizophrenia. A brain structure that is critical for temporal control is the dorsal striatum. Experience and learning can influence dorsal striatal neuronal activity, but it is unknown how these neurons change with experience in contexts which require precise temporal control of movement. We investigated this question by recording from medium-spiny neurons (MSNs) in the dorsal striatum of mice as they gained experience controlling their actions in time. We leveraged an interval timing task optimized for mice which required them to “switch” response ports after enough time had passed without receiving a reward. We report three main results. First, we found that time-related ramping activity and response-related activity increased with more experience. Second, temporal decoding by MSN ensembles improved with experience and was predominantly driven by time-related ramping activity. Finally, we found that some MSNs had differential modulation on error trials. These findings enhance our understanding of dorsal striatal temporal processing by demonstrating how MSN ensembles can evolve with experience. Our results can be linked to temporal habituation and illuminate striatal flexibility during interval timing, which may be relevant for human disease.

## Introduction

Precisely guiding movements in time is critical for human behaviors, such as cooking, driving, and crossing the street (Buhusi & Meck, 2005). Temporal control of action is disrupted in diseases such as Parkinson’s disease (PD) and schizophrenia (Ward *et al.*, 2011; Parker *et al.*, 2013; Singh *et al.*, 2021). A brain structure that is affected by both diseases and is key for controlling movements in time is the dorsal striatum (Matell & Meck, 2004; Meck, 2006). Thus, understanding how the dorsal striatum contributes to the timing of movement is important for human disease.

Dorsal striatal circuits can be explored in detail in rodent models. In rodents, disrupting striatal dopamine profoundly impairs the temporal control of action (Meck, 2006; Wang *et al.*, 2018). Striatal neurons encode time on the scale of several seconds (Matell *et al.*, 2003; Mello *et al.*, 2015; Bakhurin *et al.*, 2017; Emmons *et al.*, 2017; Wang *et al.*, 2018). For example, striatal neurons can encode temporal information by time-dependent ramping, in which firing rate changes monotonically over a temporal interval (Emmons *et al.*, 2017), although other temporal encoding schemes may also be involved (Paton and Buonomano, 2018; Zhou et. al 2020).

Task-evoked activity in striatal ensembles remaps with behavioral experience. Human brain imaging studies have found that the activity of dorsal striatal networks can be increased with sustained experience in operant tasks (Tricomi *et al.*, 2009). Moreover, dorsal striatal activity adapts with learning of specific contexts and tasks, while disrupting dorsal striatal activity can impair previously learned associations (Pasupathy & Miller, 2005; Yin *et al.*, 2005; Yin & Knowlton, 2006). The activity of striatal neurons changes markedly with sustained experience in the same procedural task even when behavior is relatively constant (Barnes *et al.*, 2005). This literature predicts that temporal encoding in the dorsal striatum should evolve with experience.

Here, we tested the hypothesis that striatal ramping changes as animals gain experience over 10 days of performing an interval timing task. We recorded from dorsal striatal ensembles of putative medium-spiny neurons (MSNs) in mice trained to perform an interval timing task which required them to “switch” from one response port to another after an interval (Balci, Ludvig, *et al.*, 2008; Balci, Papachristos, *et al.*, 2008; Tosun *et al.*, 2016). We report three main results: 1) time-related ramping patterns of activity among MSNs increased with experience; 2) temporal decoding of MSNs improved with experience; and 3) trials with erroneous responses had distinct patterns of MSN activity. Our results provide insight into how temporal encoding in striatal MSNs evolves with experience.

## Methods

### Mice

All procedures were approved by the Institutional Animal Care and Use Committee (IACUC) at the University of Iowa, and all methods were performed in accordance with the relevant guidelines and regulations (Protocol #0062039). Mice were motivated by 85% food restriction.

### Interval-timing switch task

Mice were trained to perform a switch interval timing task. Operant chambers (MedAssociates, St. Albans, VT) enclosed in sound-attenuating cabinets were equipped with two nosepokes on one wall, a food delivery port (reward hopper) located between them, a back nosepoke opposite the food delivery port, and a speaker that produced an 8 kHz tone at 72 dB. Mice were rewarded with 20-mg sucrose pellets (Bio-Serv, Flemington, NJ). First, animals were shaped to nosepoke in response to rewards using fixed-ratio task trials. These trials were initated when a mouse responded at the back nosepoke, followed by delivery of a reward after the mouse responded at an illuminated nosepoke on either side of the reward hopper. Trials were counterbalanced so that animals learned to respond at nosepokes on either the left or right side of the reward hopper. Subsequently, mice were advanced to the switch interval timing task. This task consisted of short trials (50% of trials) and long trials (50% of trials). During short trials mice were rewarded for a response at the designated short nosepoke (either the left or right nosepoke). During long trials, a response at the short nosepoke was unreinforced; instead, a response at the nosepoke on the contralateral side of the reward hopper (long nosepoke) was rewarded after 18 seconds. The nosespokes associated with short and long trials were counterbalanced to the left or right port across mice and were randomly intermixed, and the start of each trial was indicated by the same tone. Optimally, mice started by responding at the short nosepoke and switching to the long nosepoke sometime around or after six seconds (Fig 1A). The mouse’s decision to switch from the short nosepoke to the long nosepoke is a time-based decision, guided by the temporal control of action. Only long switch trials were analyzed.

**Figure 1:**
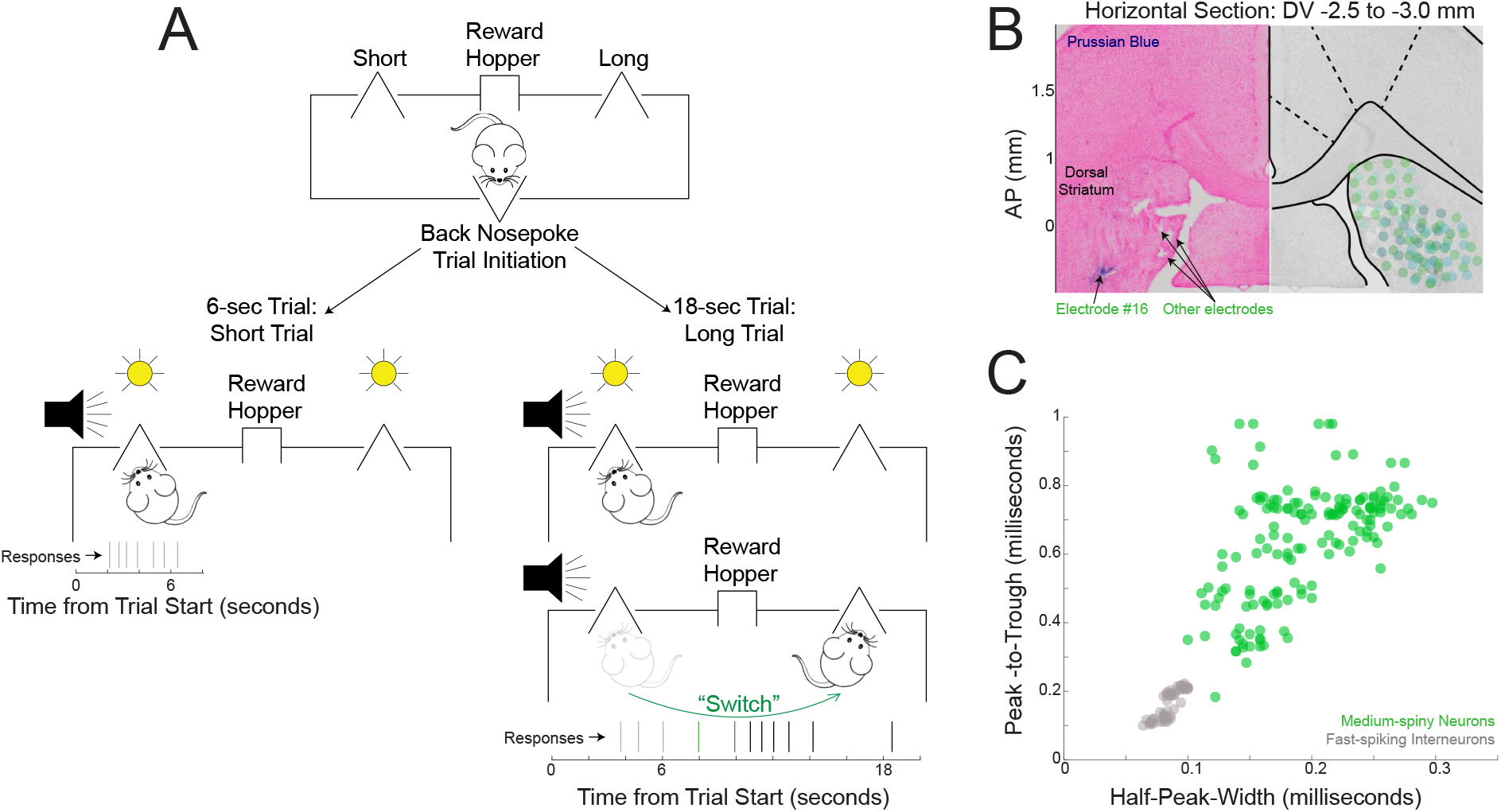
Switch interval timing task and electrode locations. A) We trained animals to perform a switch interval timing task in which mice initiate trials at a rear nosepoke. For 50% of the trials, mice were rewarded at the short nosepoke for the first nosepoke after six seconds; for the remaining trials (termed “switch trials”), mice were rewarded for the first nosepoke after 18 seconds at the long nosepoke. The temporal decision to switch from short to long nosepokes is an explicit time-based decision as in other interval timing tasks. Our analysis was focused on switch trials. B) Electrode locations in the dorsal striatum from one animal (stained with Prussian Blue on the left) and all mice (electrode locations in green on the right). C) We identified medium-spiny neurons (MSNs) based on clustering of waveform peak-to-trough distance and waveform half-peak width. Data from six mice.

Following implantation of striatal recording arrays (see below), animals were initially trained in the switch interval timing task for a period of 18 days and then acclimated to recording procedures. Recordings from the first day with >5 completed trials were considered “early training.” Mice were then trained to perform the switch interval timing task for an additional 10 days prior to recording in the “experienced” session. Mice received intraperitoneal drug injections on days 5 and 7.

### Surgical and histological procedures

C57BL/6 mice (Jackson Labs, Bar Harbor, ME) were anesthetized using IP injections of ketamine (100 mg/kg) and xylazine (10 mg/kg), and a surgical level of anesthesia was maintained using ketamine and xylazine supplements (10 mg/kg and 1 mg/kg, respectively). Craniotomies were drilled above the dorsal striatum (Fig 1B), and holes were drilled for skull screws that were connected to electrode recording arrays (MicroProbes, Gaithersburg, MD) via a separate ground wire. Microelectrode arrays were composed of 4×4 50-μm stainless steel wires (250 μm between wires and rows). These arrays were positioned in the dorsal striatum (coordinates from bregma: AP +0.9, ML ±1.4, DV −2.7 @ 0° in the posterior or lateral plane; Fig 1B) while recording neuronal activity to verify that implantation was in the correct brain area. The craniotomy was sealed with cyanoacrylate (“SloZap,” Pacer Technologies, Rancho Cucamonga, CA), and the reaction was accelerated by “ZipKicker” (Pacer Technologies) and methyl methacrylate (AM Systems, Port Angeles, WA). Mice recovered for one week before being acclimated to behavioral and recording procedures. Only the left striatum was implanted in all mice.

Following these procedures, posterolateral recording array corners (electrode #16) were marked by electrolytic lesions induced by applying a current of 50 microamps over 10 seconds through the array. After four days, mice were anesthetized using ketamine (100 mg/kg IP) and xylazine (10 mg/kg IP) and transcardially perfused with 4% formalin. Brains were post-fixed in a solution of 4% formalin and 30% sucrose, before being horizontally sectioned on a freezing microtome. Brain slices were mounted on Superfrost Plus microscope slides (Thermo Fisher Scientific, Waltham, MA) and stained for cell bodies using either cresyl violet, or nuclear fast red solution / Prussian Blue for electrode localization. Histological reconstruction was completed using postmortem analysis of electrode placement by slide-scanning light microscopy (Fig 1B; Olympus, Center Valley, PA).

### Neurophysiological recordings and neuronal analyses

Neuronal ensemble recordings were made using a multi-electrode recording system (Plexon, Dallas, TX). In each mouse, one electrode without single units was reserved for local referencing, yielding 15 electrodes per animal. After the experiments, Offline Sorter software (Plexon) was used to analyze the signals and to remove artifacts. Spike activity was analyzed for all cells that fired at rates above 0.1 Hz. Principal Component Analysis (PCA) and waveform shape were used for spike sorting. Single units were defined as those 1) having a consistent waveform shape, 2) being a separable cluster in PCA space, and 3) having a consistent refractory period of at least 2 milliseconds in interspike interval histograms. Putative MSNs were further separated from striatal interneurons based on hierarchical clustering of the waveform peak-to-trough ratio and the half-peak width (*fitgmdist* and *cluster.m*; Fig 1C) (Berke, 2011). All neuronal analyses focused on putative MSNs. Note that the same electrodes are recorded from across sessions.

### Statistics

All data and statistical approaches were reviewed by the Biostatistics, Epidemiology, and Research Design Core (BERD) at the Institute for Clinical and Translational Sciences (ICTS) at the University of Iowa. All code and data are available at http://narayanan.lab.uiowa.edu. For behavioral data, we used linear mixed-effects models where the outcome variable was response time on long trials and the predictor variable was session type (“early training” vs. “experienced”); animals were included as a random effect. We used the median to measure central tendency and the interquartile range to measure variance across mice. Wilcoxon nonparametric signed-rank or rank-sum tests were used to compare behavior between early training and experienced sessions. Wilxocon signed-rank tests were also used to compare decoding analyses between early training and experienced sessions and different neuronal ensembles. To quantify effect size, we used Cohen’s d. To present evidence related to the null hypothesis, we calculated the Bayes Factor (Morey & Rouder, 2011).

Analyses of neuronal activity and basic firing properties were carried out using NeuroExplorer (Nex Technologies, Littleton, MA) and custom routines for MATLAB (Parker *et al.*, 2014; Emmons *et al.*, 2017, 2019). Neuronal modulations were quantified using generalized linear models (GLMs) at the individual neuron level. For each neuron, we constructed a model in which the response variable was the firing rate binned at 0.2 seconds and the predictor variable was either time in the interval or motor response. All models were run at a trial-by-trial level to avoid effects of trial-averaging (Latimer *et al.*, 2015). For each neuron, a p-value was obtained for main effects of time in the interval (i.e., “time-related ramping”) or response. Across the ensemble, p-values were corrected via Benjamini-Hochberg false-discovery-rate (FDR), with values <0.05 considered significant for each neuron. To identify neurons with different activity at short vs. long or right vs. left nosepokes, we compared firing rates in the 0.2 seconds around nosepoke response via a Wilcoxon rank-sum test for each neuron. We compared GLM-defined neuronal modulations between early training and experienced sessions via a X^2^ test, assuming neurons were independent as in our past work (Narayanan & Laubach, 2006; Parker *et al.*, 2014, 2015; Emmons *et al.*, 2017; Kim *et al.*, 2017). For comparisons of ramping-related or response-related modulations between early training and experienced sessions, we used logistic regression and included a random intercept for each animal to control for mouse-specific effects. For error-related modulations, a similar approach was taken, but the GLM was constructed where the reponse variable was firing rate, and the predictor variable was whether the trial was correct or had erroneous nosepokes; only neurons from animals with >5 trials of each type were analyzed. A p-value was obtained for main effects of errors for each neuron and corrected via FDR across the ensemble.

We used a naïve Bayesian classifier to examine neuronal ensemble decoding, as we have in our past work (Emmons *et al.*, 2017; Kim *et al.*, 2017). We calculated kernel density estimates (bandwidth: 0.5) of trial-by-trial MSN firing rates, binned at 1 second, from all neurons with at least eight trials. To prevent edge effects that might bias classifier performance, we included data from six seconds prior to trial start and six seconds after interval end. We used leave-one-out cross-validation to predict an objective time from firing rate within a trial. We evaluated classifier performance by computing the R^2^ of objective time vs. predicted time, only for bins during the interval. With perfect classification, the R^2^ would approach 1. Classifier performance was compared to ensembles with time-shuffled firing rates via a Wilcoxon signed-rank test.

## Results

Our goal was to study how striatal temporal encoding evolves with experience. We tested this idea by implanting multielectrode recording arrays targeting the dorsal striatum and training six animals in the switch interval timing task for ~10 days (Fig 1A). For animals in early training, the average short nosepoke response time was 9.3±2.8 seconds (median±interquartile range) and 13.7±3.3 seconds at the long nosepoke. For experienced animals, the average response times were 7.5±1.4 seconds at the short nosepoke and 14.8±1.2 seconds at the long nosepoke. Mice did not reliably perform more trials in experienced sessions compared to early training (31±26 trials in early training vs. 39±21 trials in experienced sessions, Wilcoxon signed-rank p=0.13, Bayes factor=0.74). A linear mixed-effects model revealed that there was a highly-significant main effect of short vs. long nosepoke (F(1,2788)=27.7, p<0.05), a main effect of experience (F(1,2788)=7.5, p=0.006), and a significant interaction (interaction: F(1,2788)=3.9, p=0.05; model fit R^2^=0.20; Fig 2A-E). Although there was no consistent change in switch times (9.3±3.6 seconds vs. 9.3±1.5 seconds; Wilcoxon signed-rank p=0.8; Bayes factor=2.5; Fig 2F) or the coefficient in variation (36.3±17.9% vs. 34.5±20.8%; Fig 2G), mice performed better with experience, with a higher percentage of trials terminated by a response at the correct response port (naïve: 59±22% vs. experienced 80±23%; Wilcoxon signed-rank p=0.03; Cohen’s d=1.3; Fig 2H).

**Figure 2:**
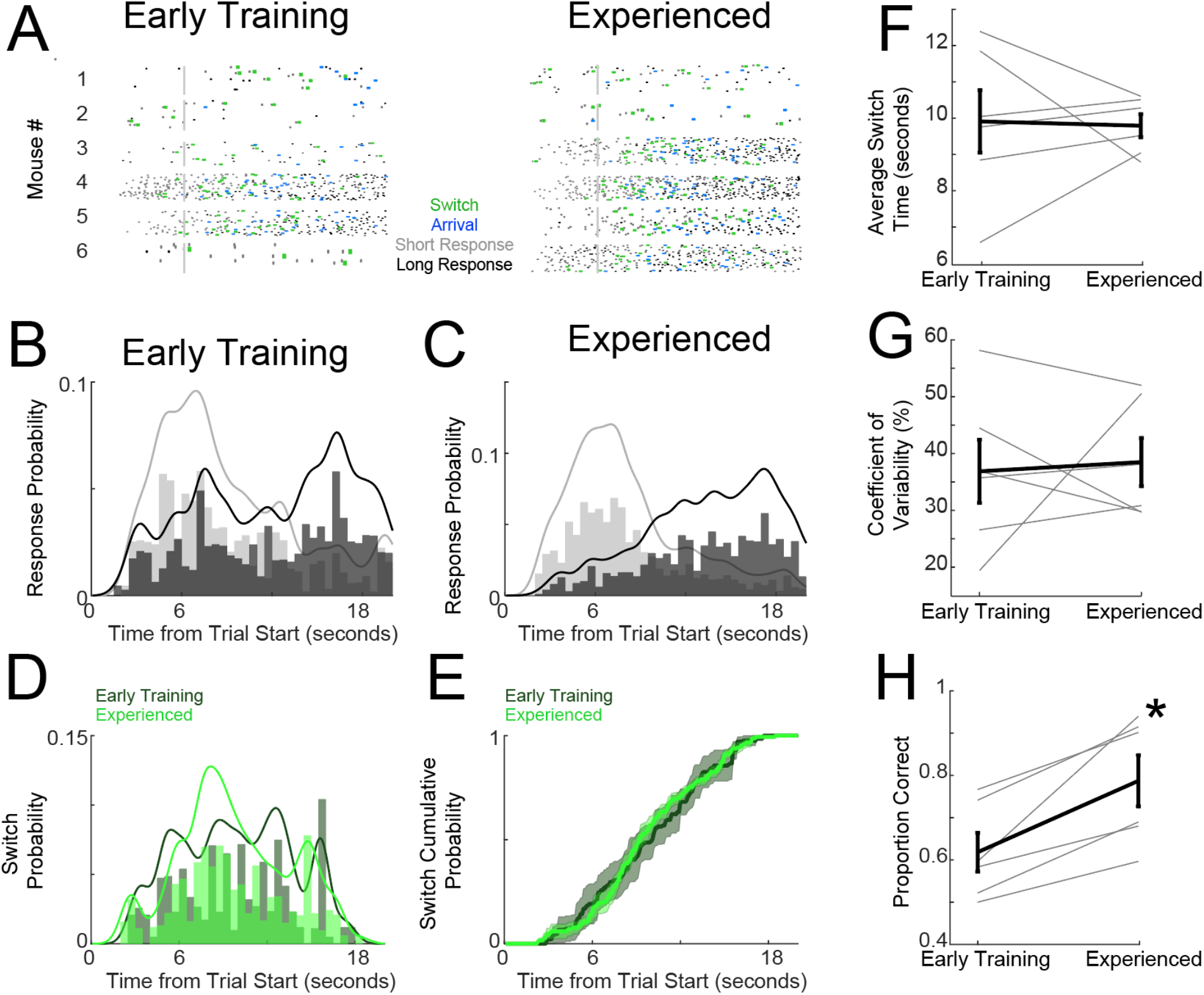
Interval timing performance improves with experience. Rasters of short nosepokes (gray), long nosepokes (black), and switch nosepokes (green) in (A) early training vs. experienced animals. B) Time-response histograms of short (gray) vs. long (black) nosepokes for early training vs. C) experienced animals. D) Distribution or E) cumulative probability of switch responses (when animals departed from the short nosepoke and subsequently responded at the long nosepoke) did not change with experience. F) Mean switch times or G) the coefficient of variability did not change with experience. H) The proportion of correct responses increased with experience. Gray lines in F-H are individual animals; black lines are mean±SEM.*=p<0.05 via Wilcoxon signed-rank. Data from 445 trials in six animals.

We were interested in medium spiny neuron (MSN) activity during the switch interval-timing task. We recorded from 77 MSNs in early training and 79 MSNs in the same animals that had become experienced after extensive training. To quantify experience effects at a trial-by-trial level, we turned to GLMs in which the outcome variable was firing rate and the predictor variable was either time in the interval or motor response. Neurons with an FDR-corrected significant main effect of time were considered to exhibit time-related ramping activity (Fig 3A), and neurons with FDR-corrected significant main effects of response were considered response-related (Fig 3B). Additionally, many neurons displayed both time-dependent ramping and response-related changes in their activity (Fig 3C). We found that the total number of time-related ramping neurons increased with experience (29% vs. 44%, X^2^_(1,156)_=4.2, p=0.04; p=0.01 via mixed-effect logistic regression; Fig 3C-D), as did the total number of response-related neurons (28% vs. 46%; X^2(1,156)^ =5.6, p=0.02; p=0.01 via mixed-effect logistic regression; Fig 3C-D). Additionally, the number of neurons that showed both time-related ramping and response-related modulation increased with experience (10% vs. 25%; X^2^_(1,156)_=6.0, p=0.01; Fig 3C-D). Of note, there were response-related neurons that were modulated by short vs. long responses, but these did not significantly change with experience (14 of 21 response-related neurons in early training and 16 of 36 neurons in experienced sessions had short vs. long activity; X^2^_(1,57)_=2.6, p=0.11). We found that 9% of neurons in early training sessions were modulated by the switch response, when mice transitioned from short to long nosepokes, and 10% were modulated in experienced sessions (X^2^_(1, 156)_=0.2, p=0.66). These data suggest that ramping increased with experience, both on its own and where it interacted with response-related activity.

**Figure 3:**
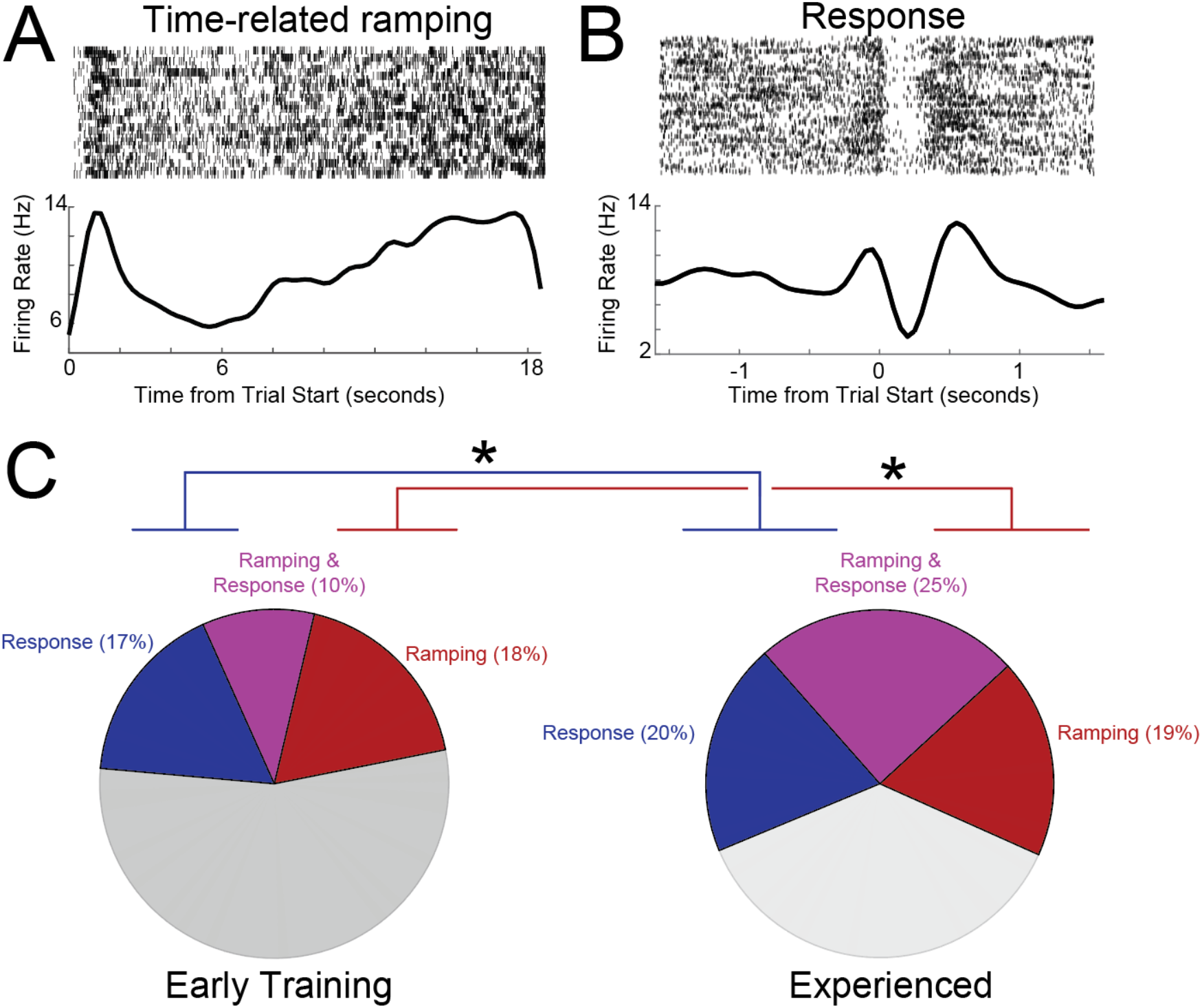
Response-related and timing-related MSNs increase with experience. A) Example of an MSN with time-related ramping activity. B) Example of an MSN with response-related activity. C) Pie charts illustrating the frequency of MSNs exhibiting significant response-related activity (blue), time-related ramping activity (red), or both (purple), during interval timing in early training vs. experienced sessions. * by ramping- and response-related activity indicates that they both increased in experienced sessions with p<0.05 as calculated by X^2^-test. Data from 80 MSNs in six mice during early training sessions and 81 MSNs in the same six mice during experienced sessions.

To examine the computational consequences of this shift, we turned to decoding analyses where we used naïve Bayesian classification to predict time from striatal ensembles (Emmons et al., 2017; Kim et al., 2017). Consistent with the change in time-related ramping with experience, we found that experienced striatal ensembles predicted time with greater accuracy than in early training (R^2^: early training: 0.29±0.41 vs. experienced: 0.62±0.60; Wilcoxon signed-rank p=0.02; Cohen’s d=0.8; Fig 4A-C). Of note, both ensembles were reliably more predictive of time than shuffled data (early training shuffled R^2^: 0.01±0.04; Wilcoxon signed-rank vs. early training p=0.0002; experienced shuffled R^2^: 0.11±0.18; Wilcoxon signed-rank vs. early training p=0.0005). In experienced sessions we found that MSNs with both time-related ramping and response-related activity were similarly predictive of time (ramping R^2^: 0.38±0.38; ramping & response R^2^: 0.47±0.35; Wilcoxon signed-rank p=0.28; Bayes factor=3.0; data from experienced sessions only; Fig. 4C). Critically, this was stronger than MSNs with response-related activity alone (0.10±0.31; Wilcoxon signed-rank p=0.001 vs. ramping; Cohen’s d=1.2; p=0.007 vs. ramping&response:; Cohen’s d=0.9; Fig. 4D). Furthermore, striatal ensembles without time-related ramping did not decode time well (0.14±0.18; Wilcoxon signed-rank p=0.0003 vs. ramping; Cohen’s d=1.4; p=0.002 vs. time-related ramping&response; Cohen’s d=1.1; Fig. 4D). These data make three key points: 1) ensembles with time-related ramping contributed strongly to temporal decoding, whether or not they had response-related activity, 2) ensembles without time-related ramping had poor temporal decoding, and 3) response-related neuronal activity, despite increasing with experience, did not decode time. These data are consistent with previous findings from our group in that neurons with time-related ramping strongly convey temporal information (Emmons et. al., 2017) and that temporal decoding improves in striatal ensembles with experience (Emmons et. al., 2020).

**Figure 4:**
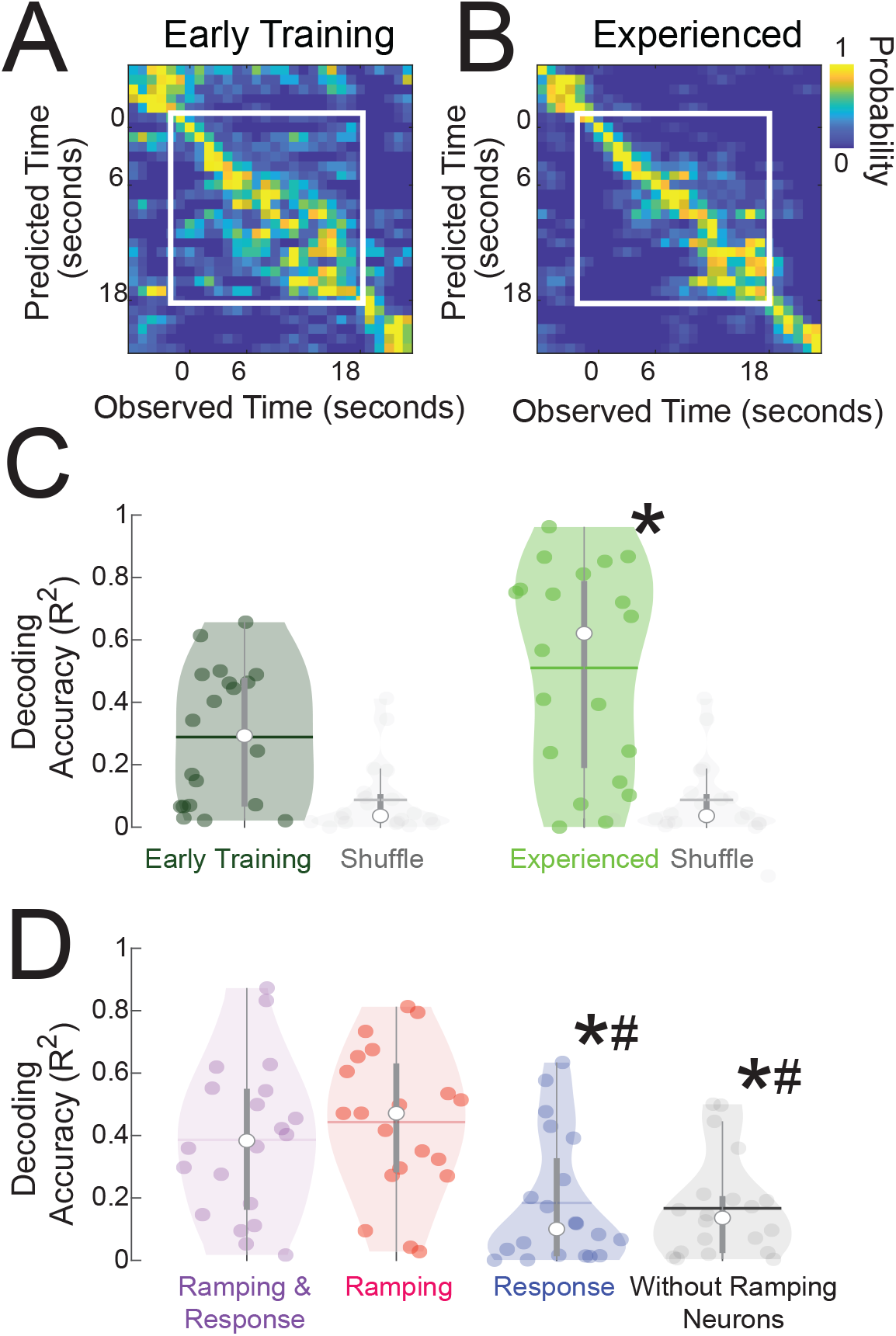
MSN ensembles improve temporal decoding with experience. We trained naïve Bayesian classifiers to predict time from firing rate on a trial-by-trial basis. Temporal predictions for A) early training and B) experienced sessions; predicted time is on the y-axis and observed time is on the x-axis, with yellow representing the highest probability. Only the time during the trial (0–18 seconds; white box) is analyzed. C) Temporal decoding improved for early training vs. experienced sessions and was consisently stronger than shuffled data. *=p<0.05 via Wilcoxon signed-rank. D) Temporal decoding was higher for MSNs with both time-related and response-related modulations than for MSNs with response-related modulations. *=p<0.05 via Wilcoxon signed-rank vs. ramping & response neurons, and #=p<0.05 via Wilcoxon signed-rank vs. ramping neurons. Critically, MSNs with only response-related modulations were significantly worse for classification than MSNs with ramping or ramping & response-related modulations; without ramping neurons, temporal decording accuracy decreased. Data from MSN ensembles in six animals during early training and experienced sessions.

Finally, we examined neuronal activity on error trials in which animals did not successfully switch from short to long nosepokes (Fig 5). We only examined neurons that had five or more errror trials; we recorded 61 neurons in early training and 37 neurons in experienced sessions among our six mice. In early training, 51% of these neurons were error-modulated, which was not consistently different from experienced sessions (35%, X^2^_(1,98)_=2.8, p=0.10; Fig 5B). These data indicate that error trials could involve distinct patterns of striatal activity but that these patterns did not consistently incfease with experience. Together, these results together describe how ramping modulations, response modulations, and temporal decoding increase with experience while error-related activity does not.

**Figure 5:**
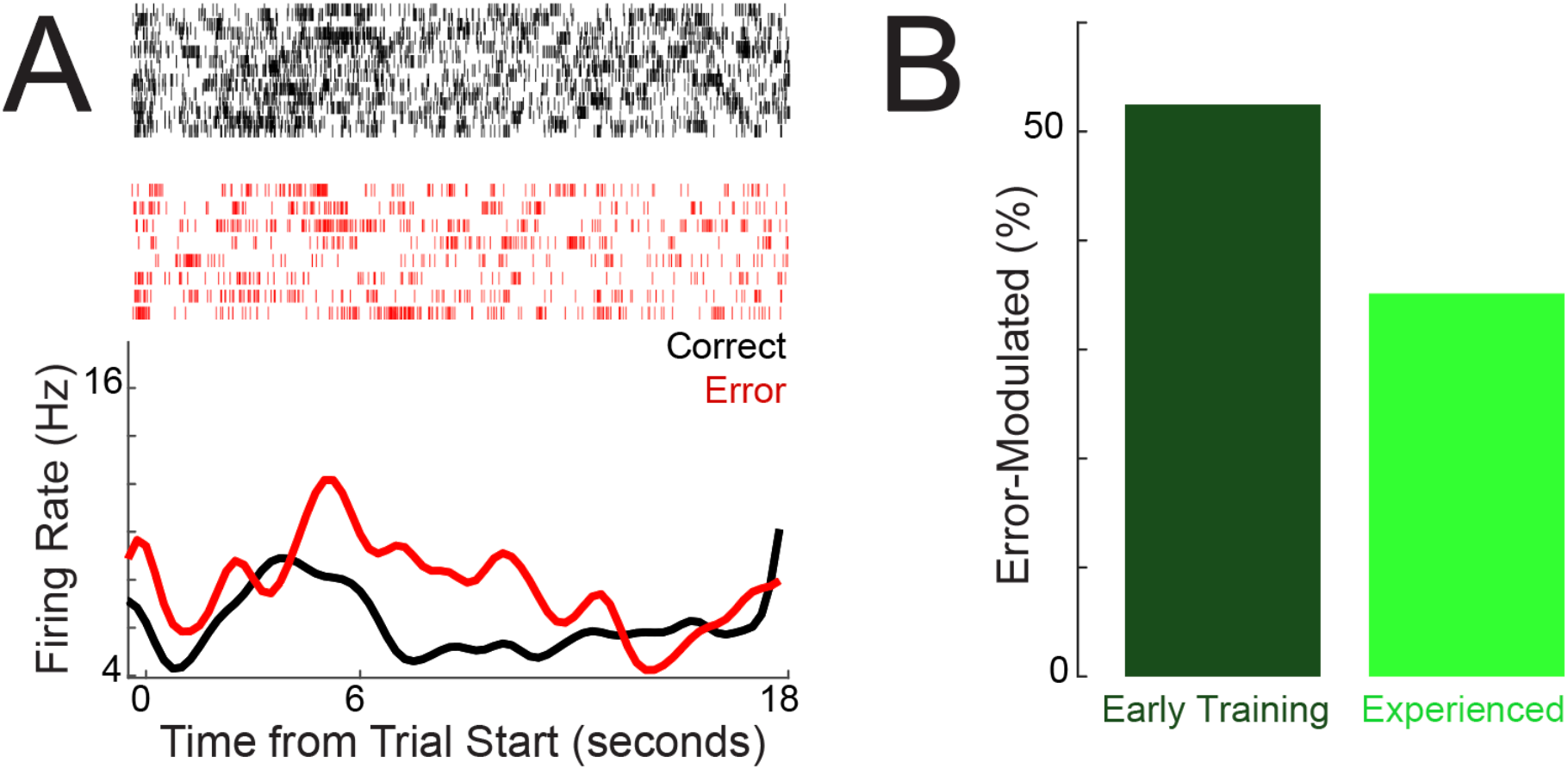
Neuronal activity on error trials. A) An MSN with distinct activity during error-related trials. B) Fraction of error-trial modulated neurons in early training vs. experienced sessions. Data from 98 neurons with >5 error trials in six mice during early training and experienced sessions.

## Discussion

We explored how striatal temporal encoding remapped with experience. First, we found that patterns of MSN activity evolved after 10 days of interval timing. Second, we found that dorsal striatal ensembles improved temporal decoding with experience. This was driven largely by MSNs with time-related ramping activity, as MSNs without this activity had poor temporal decoding. Finally, we found that dorsal striatal neurons could have distinct patterns of error-related activity. Taken together, these data provide evidence that MSN temporal encoding can evolve with experience.

These data are broadly consistent with our previous work (Emmons *et al.*, 2016, 2017, 2019, 2020). In one of these studies with well-trained animals (Emmons *et al.*, 2020), we found that striatal time-related ramping was stable as highly-trained rats learned a new, shorter fixed interval over three days. By contrast, in this study we compared striatal temporal encoding just after mice aquired a switch interval timing task and again after ~10 days of experience. We note that striatal decoding improved in both studies, providing convergent evidence that temporal encoding by striatal ensembles is enhanced with experience.

Movement confounds studies of the temporal control of action (Namboodiri & Hussain Shuler, 2014; Paton & Buonomano, 2018). We have previously controlled for motor confounds via regression (Parker *et al.*, 2014; Kim *et al.*, 2017; Emmons *et al.*, 2019), and we note that time-related ramping is still observed prior to the first response (Emmons *et al.*, 2017). In switch interval timing, response rates at the short interval do not monotonically increase with time; indeed response rates at the long interval increase over the interval but plateau in experienced animals, and switch interval times are mostly distributed after six seconds (Fig 2D); Balci, Papachristos, *et al.*, 2008). Supporting this idea are our observations that response-related activity alone did not decode time (temporal decoding was driven by time-related ramping activity), and that neurons multiplexing time and movement could change with experience. Finally, time-related ramping can reflect averaging across a session (Latimer *et al.*, 2015). For this reason, our definition of time-related ramping was derived from trial-by-trial GLMs that quantify trial-by-trial main effects of firing rate over the interval (Emmons *et al.*, 2017), although additional analytic tools may help further resolve these dynamics (Narayanan, 2016; Zylberberg & Shadlen, 2016).

There are many alternative coding schemes that robustly represent temporal information, including those based on oscillatory activity, sequences, and network states (Paton & Buonomano, 2018; Zhou *et al.*, 2020). While those features may be latent within our dorsal striatal ensembles, we have reported consistent evidence that time-related ramping contributes to temporal decoding (Narayanan & Laubach, 2009; Parker *et al.*, 2014; Narayanan, 2016; Emmons *et al.*, 2017; Kim *et al.*, 2017). We have also shown that striatal ramping is correlated with prefrontal ramping and requires prefrontal top-down control (Emmons *et al.*, 2017, 2019). While other coding schemes may be relevant, we found that striatal ensembles without ramping activity decode time poorly. Taken together, these data suggest that time-related ramping contributes to temporal processing in tasks that require temporal control of action.

We report that striatal neurons exhibit different activity profiles on error trials in which animals failed to switch from the short to the long nosepoke. This analysis was not possible in our previous work with the fixed-interval timing task or in many prior studies of interval timing in which all trials with responses after a target interval were rewarded (Matell *et al.*, 2003; Parker *et al.*, 2014, 2015; Bakhurin *et al.*, 2017; Emmons *et al.*, 2017, 2020). In addition, despite increases in time-related ramping, response-related activity and fewer errors with experience, error-related activity did not change with experience. This finding suggests that error-related firing is not strictly related to time-related ramping or responding; future studies may further elucidate these error-related patterns, particulary in contexts when mice make enough errors to facilitate robust analyses.

Finally, these data illuminate the neuronal basis of temporal habits. Indeed, the dorsal striatum is critical for habits in which stimulus-response associations persist in overtrained animals, despite changes in reward contingencies (Yin *et al.*, 2005; Yin & Knowlton, 2006; Balleine *et al.*, 2007). Accordingly, past work has found that the striatum profoundly remaps with overtraining (Barnes *et al.*, 2005). We did not manipulate rewards in the present study, as we found relatively sparse reward representations in our striatal ensembles. However, we found profound remapping in striatal ensemble encoding of movements and time, which may represent the neural basis of temporal habits. This idea is supported by our behavioral data suggesting that switch time and variation do not consistently change with experience. Future studies will explore how reward devaluation might influence interval timing and other stimulus-response associations separated by time.

Dorsal striatal neurons encode many aspects of behavior (Kimchi *et al.*, 2009; Kimchi & Laubach, 2009). We report that time-related ramping increases with experience during interval timing. This knowledge is germane to human diseases that affect the striatum such as Parkinson’s disease (PD). Indeed, PD patients can have specific challenges in learning (Frank *et al.*, 2004; Nieuwboer *et al.*, 2009) and in performing functions for which they have extensive experience (Uc *et al.*, 2006, 2007). Understanding the striatal basis of temporal encoding might lead to new treatments for human diseases such as Parkinson’s disease and schizophrenia that degrade learning, cognition, and previously-formed habits.

Our work has several limitations. The first is that we cannot dissociate MSN subtypes that express D1 vs. D2 dopamine receptors, and we are uncertain how the powerful neuromodulator dopamine affects striatal networks (Cui *et al.*, 2013). Second, we cannot fully dissociate ramping activity from movements, which would require alternative task designs (Church & Deluty, 1977), although one critical advantage of our task design is that it readily translates to humans (Ward *et al.*, 2011; Kim *et al.*, 2017; Parker *et al.*, 2017), particularly with Parkinson’s disease (Singh *et al.*, 2021). Third, with our approach, we were unable to identify oscillatory or state-based coding schemes in the striatum that change with experience, although our analytical approach may not be suited to identify these network properties.

Despite these limitations, the present study provides evidence that temporal encoding in the dorsal striatum changes with experience. Specifically, we find that time-related ramping can increase, resulting in improved temporal decoding. Our results illustrate how striatal ensembles remap with temporal experience.

## Acknowledgements

This work was funded by NIH MH116043 to NSN. This study was supported in part by the University of Iowa Institute for Clinical and Translational Science, which is granted with Clinical and Translational Science Award funds from the National Institutes of Health (UL1TR002537).

## Author contributions

RAB and NSN designed the experiments. RAB collected the data, with assistance from RAV, MO, and MAW. RAB and YK wrote the code. EE and YK checked the analysis and code. RAB, MAW, and NSN wrote the paper.

